# A novel statistical framework for meta-analysis of total mediation effect with high-dimensional omics mediators in large-scale genomic consortia

**DOI:** 10.1101/2024.04.29.591700

**Authors:** Zhichao Xu, Peng Wei

**Affiliations:** Department of Biostatistics, The University of Texas MD Anderson Cancer Center

## Abstract

Meta-analysis is used to aggregate the effects of interest across multiple studies, while its methodology is largely underexplored in mediation analysis, particularly in estimating the total mediation effect of high-dimensional omics mediators. Large-scale genomic consortia, such as the Trans-Omics for Precision Medicine (TOPMed) program, comprise multiple cohorts with diverse technologies to elucidate the genetic architecture and biological mechanisms underlying complex human traits and diseases. Leveraging the recent established asymptotic standard error of the R-squared (*R*^2^)-based mediation effect estimation for high-dimensional omics mediators, we have developed a novel meta-analysis framework requiring only summary statistics and allowing inter-study heterogeneity. Whereas the proposed meta-analysis can uniquely evaluate and account for potential effect heterogeneity across studies due to, for example, varying genomic profiling platforms, our extensive simulations showed that the developed method was more computationally efficient and yielded satisfactory operating characteristics comparable to analysis of the pooled individual-level data when there was no inter-study heterogeneity. We applied the developed method to 8 TOPMed studies with over 5800 participants to estimate the mediation effects of gene expression on age-related variation in systolic blood pressure and sex-related variation in high-density lipoprotein (HDL) cholesterol. The proposed method is available in R package `MetaR2M` on `GitHub`.

## 1 Introduction

Large-scale genomic consortia and biobanks have facilitated genetic and genomic research by providing data and tools to probe into complex human diseases and traits with unparalleled depth and applicability (Fatumo et al., 2022; Wang et al., 2022; Bycroft et al., 2018). For instance, in our motivating example, the National Heart, Lung, and Blood Institute’s (NHLBI) Trans-Omics for Precision Medicine (TOPMed) project brings together over 85 cohorts consisting of more than 180,000 participants using various high-throughput profiling technologies to elucidate the genetic architecture and biological mechanisms underlying complex human traits (Taliun et al., 2021). Advances in technology and data sharing have made individual participant data more accessible (Chalmers, 1993; Sutton et al., 2000). However, the acquisition and analysis of such individual-level data is time-consuming, financially demanding, and limited by privacy concerns.

High-dimensional mediation analysis is a crucial analytical approach focused on evaluating the mediating role of molecular phenotypes, such as gene expression, in the relationship between environmental exposure/risk factor and health outcomes (VanderWeele, 2016; Vo et al., 2020; Dai et al., 2022; Derkach et al., 2020; Zhang et al., 2018; Zeng et al., 2021). A variance-based R-squared measure, denoted as 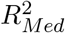, was proposed to estimate the total mediation effect in the high-dimensional setting (Yang et al., 2021; Chi et al., 2024). A recently developed two-stage cross-fitted interval estimation procedure for 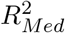 enables the implementation of meta-analysis in mediation analysis due to its availability of asymptotic standard error and computational efficiency (Xu et al., 2023), as to be pursued here.

Meta-analysis is a powerful tool for synthesizing the effects of interest across multiple similar individual studies (Borenstein et al., 2021; Brockwell and Gordon, 2001). Established meta-analysis techniques use summary statistics to resolve the difficulties in accessing individual-level data (DerSimonian and Laird, 1986; Mantel and Haenszel, 1959; Lu and Ades, 2004; Li et al., 2023; Zhu et al., 2015). Fixed-effects meta-analysis stands out as the most widely-used and robust method for combining findings from multiple genetic studies (Pfeiffer et al., 2009). Fixed-effects models require the assumption that the true effects of interest are identical across all studies. Within this domain, the inverse variance weighting method is widely adopted, attributing weights to each study based on the inverse of the sampling variance of the estimator of interest, for example, estimated odds ratio for binary data (Kavvoura and Ioannidis, 2008). The Mantel-Haenszel method computes a weighted average of odds ratios, with weights being proportional to the size and variability of each study (Mantel and Haenszel, 1959). Random-effects models are used when there is heterogeneity across the studies in the meta-analysis. The DerSimonian and Laird (DL) estimator is favored for its simplicity and robustness (DerSimonian and Laird, 1986). Several authors have highlighted the importance of including a considerable number of studies in the random-effects meta-analysis to ensure the reliability of inferential results(Van Houwelingen et al., 2002; Guolo, 2012). More recently, the median-unbiased Paule-Mandel (MPM) estimator has been proposed to estimate the heterogeneity from the median of the generalized *Q* statistic proposed by Cochran instead of its expected value (Cochran, 1954; Viechtbauer, 2021).

Meta-analysis and systematic reviews have been extensively applied in mediation analysis to identify potential mediators influencing health-related outcomes (Gu et al., 2015; Lubans et al., 2008; Lee et al., 2015; Mansell et al., 2013; Satten et al., 2022). However, its methodology is largely underexplored in high-dimensional mediation analysis, particularly in estimating the total mediation effect of high-dimensional omics mediators (Zeng et al., 2021). To this end we introduce a novel meta-analysis framework, allowing for both fixed-effects and random-effects, to estimate the total mediation effect in high-dimensional settings. This framework requires only summary statistics and allows between-study heterogeneity arising from factors such as differences in high-throughput technologies (microarray v.s. RNA-sequencing) and diverse ethnicity. Our extensive simulations show that the efficiency and coverage probability when using summary statistics are comparable to those achieved with the individual-level data in meta-analysis. Applying this innovative framework, we conducted a meta-analysis across various cohorts from the TOPMed Framingham Heart Study (FHS) and the Multi-Ethnic Study of Atherosclerosis (MESA) to estimate the mediation effects of gene expression on age-related variation in systolic blood pressure (BP) and sex-related variation in high-density lipoprotein (HDL) cholesterol. The proposed meta-analysis framework is implemented in the R package `MetaR2M` available on `GitHub` and to be submitted to R/CRAN.

## 2 Methods

In this section, we provide the background of mediation models, potential mediators/non-mediators, and the *R*^2^-based total mediation effect. Then we review the fixed-effects and random-effects models in meta-analysis using summary statistics versus individual-level data, followed by the proposed framework for meta-analysis of total mediation effect under high-dimensional settings.

### 2.1 Mediation models and 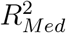 measure

Let *X* denote a *n ×* 1 vector of the exposure variable, ***M*** denote a *n × p* matrix for *p* potential mediators, *M*_*j*_ be a *n ×* 1 vector for the *j*th mediator, and *Y* represent a *n ×* 1 vector of the outcome variable. Without loss of generality, we assume that all variables have been centered at 0 and scaled to have variance of 1; in addition, all measured potential confounders have been regressed out from *X, M*_*j*_’s and *Y* from the following equations, which constitute the mediation model:

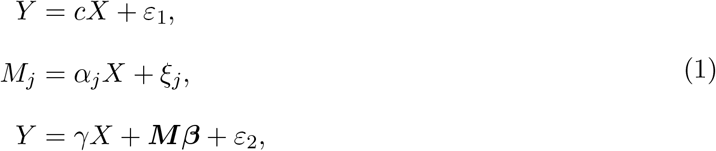

where *c*, ***α*** = (*α*_1_, …, *α*_*p*_)^*′*^, ***β*** = (*β*_1_, …, *β*_*p*_)^*′*^, and *γ* are the coefficients of regressions that can be estimated via maximum likelihood estimation (MLE), and *ε*_1_, *ε*_2_, and *ξ*_*j*_ = (*ξ*_1*j*_, …, *ξ*_*nj*_)^*′*^ are *n ×* 1 vectors of random errors. Here parameter *c* is the total mediation effect linking the exposure to the outcome, and *γ* captures the direct effect of *X* on *Y* in the classical mediation analysis framework. As illustrated in Fig 1, we categorize the potential mediators 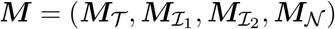 into four groups: true mediators and three types of non-mediators (Baron and Kenny, 1986). True mediators ***M***_*𝒯*_ (Fig 1A) are the variables associated with both the exposure and the outcome (*α*_*j*_ *≠*0, *β≠* 0 for *j ∈ 𝒯*). Non-mediators 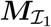(Fig 1B) are the variables associated with the outcome but not the exposure (*α*_*j*_ = 0, *β*_*j*_ *≠*0 for *j ∈ ℐ*_1_). Similarly, non-mediators 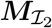(Fig 1C) are the variables associated with the exposure but not the outcome (*α*_*j*_*≠* 0, *β*_*j*_ = 0 for *j ∈ ℐ*_2_). Lastly, non-mediator noise variables ***M***_*𝒩*_ (Fig 1D) are the variables not associated with either the exposure or the outcome (*α*_*j*_ = *β*_*j*_ = 0 for *j ∈ 𝒩*). The inclusion of the specific type o f n on-mediators i n the high-dimensional mediation analysis could potentially bias the estimation (Yang et al., 2021).

**Figure 1:**
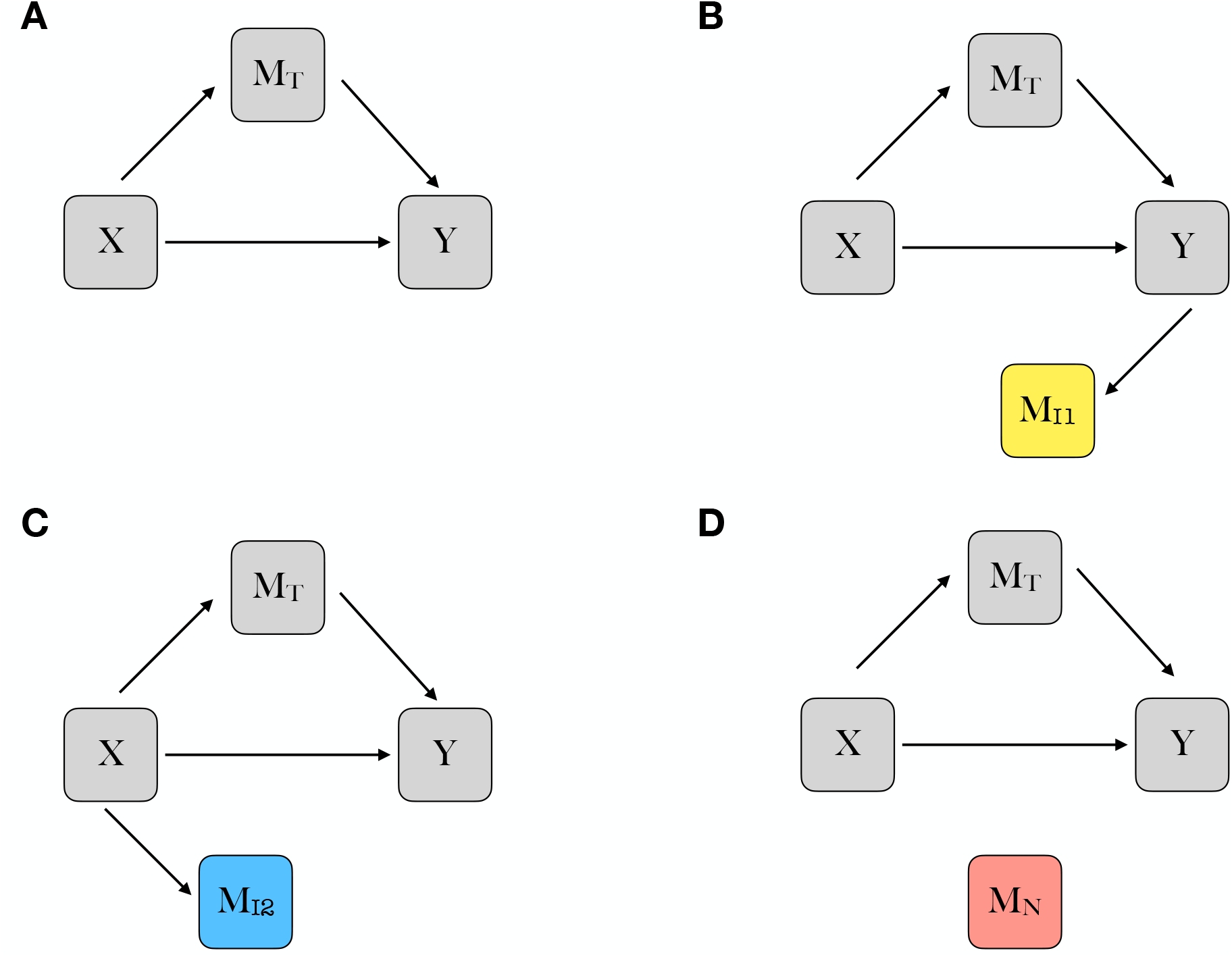
Graph representations of potential mediation models. X refers to the exposure variable. Y refers to the outcome variable. ***M***_*T*_ refers to the true mediators associated with both the exposure and the outcome. 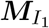 refers to the non-mediators associated with the outcome but not the exposure. 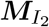 refers to the non-mediators associated with the exposure but not the outcome. ***M***_*N*_ refers to the non-mediators noise variables that are not associated with either the exposure or the outcome.

From Eq (1), the second-moment-based total mediation effect measure 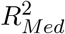 in the multiplemediator setting is defined as follows:

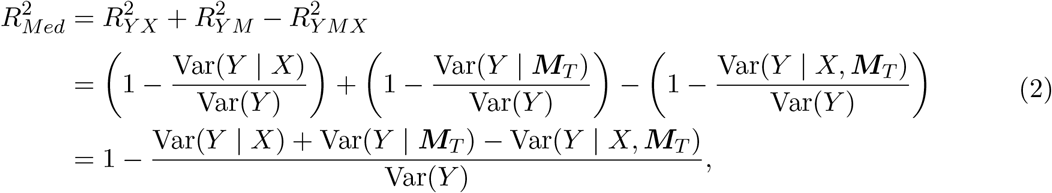

where 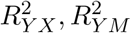 and 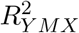 represents the coefficient of determination for the regression models in which *Y* is regressed on *X*, ***M***_*𝒯*_, and (*X*, ***M***_*𝒯*_), respectively. 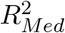 as a measure of total mediation effect is interpreted as the amount of variation in the outcome *Y* that is explained by exposure *X* through mediators *M* (Yang et al., 2021; Chi et al., 2024). Note that when *X, ξ*_*j*_, *ε*_1_, and *ε*_2_ are independently distributed, Eq (2) remains valid if we substitute ***M***_*𝒯*_ with ***M***_*𝒮*_ where *𝒮* is the union of the true mediators ***M***_*T*_ and the non-mediators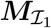, denoted as *𝒮* = *𝒯 ∪ ℐ*_1_ (Xu et al., 2023).

Using 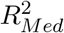 as a measure of the total mediation effect offers several advantages. First, 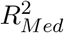 is an appealing complementary measure to traditional total mediation effect measures, such as the product measure for mediation/indirect effect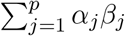, by avoiding the issue of cancellation from component-wise mediation effects *α*_*j*_*β*_*j*_’s of different directions (VanderWeele, 2015; Judd and Kenny, 1981). Second, since 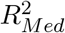 is defined based on the coefficient of determination *R*^2^, it allows the mediators to be correlated which is likely the case in high-dimensional genomics settings (Yang et al., 2021). Third, 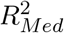 can be extended beyond continuous outcomes, such as time-to-event outcomes, which relax the rare event assumption as required by the product measure (Chi et al., 2024).

### 2.2 Meta-analysis framework for 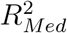measure

Recently, a novel two-stage interval estimation procedure using cross-fitted Ordinary Least Squares (OLS) regressions, CF-OLS, for estimating 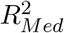 in a single study has been proposed (Xu et al., 2023). This method is based on cross-fitting and sample-splitting techniques and is tailored for estimating the confidence interval of total mediation effect in high-dimensional mediators settings (Xu et al., 2023). The newly derived asymptotic distribution and, thus, the standard error, of the estimator makes it possible for meta-analysis using summary statistics, i.e., point estimate and standard error of the estimated total mediation effect from each study.

In CF-OLS, following the data split into two subsamples, the initial step involves variable selection. It is worth noting that the presence of non-mediator 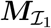 and noise variables ***M***_*𝒩*_ does not affect the estimation when all true mediators and non-mediators are independent. However, non-mediator 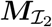 can introduce bias and inconsistency, especially in high-dimensional settings (Yang et al., 2021). Therefore, we used the iterative Sure Independence Screening (iSIS) (Fan and Lv, 2008) in conjunction with the Minimax Concave Penalty (MCP) (Zhang, 2010) screening procedure, known as iSIS-MCP, to identify and filter out the non-mediator 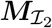. Subsequently, we applied the False Discovery Rate (FDR) procedure to further exclude non-mediator 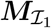 and noise variables ***M***_*𝒩*_, as they might bias the results when they are highly correlated (Xu et al., 2023). With the true mediators ***M***_*𝒯*_ selected in each of the two subsamples, the inference of 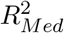 is conducted based on the asymptotic standard error of its estimator 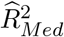. After the variable selection procedure, we will have

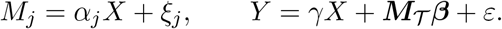

If certain assumptions are met and the mediator selection satisfies the sure screening property (Xu et al., 2023), then it holds that

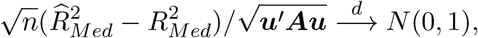

where ***u*** = (1*/* Var(*Y*), −1*/* Var(*Y*), −1*/* Var(*Y*), (Var(*Y* | *X*)+Var(*Y* | ***M***_*𝒯*_)−Var(*Y* | *X*, ***M***_*𝒯*_))*/* Var(*Y*)^2^)^*′*^ and ***A*** is the (constant) covariance matrix of (*ε*^2^, *η*^2^, *ζ*^2^, *Y* ^2^) (Xu et al., 2023). Specifically,

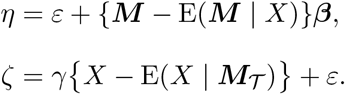

The above result indicates that 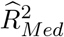 is a consistent estimator of 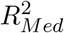 and follows a normal distribution, based on which the standard error and a 95% confidence interval can be analytically derived. We estimate the asymptotic covariance matrix ***A*** by the residuals of the corresponding linear regressions via the OLS.

Fig 2 illustrates the workflow of our proposed meta-analysis approach to estimating 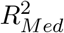 from

**Figure 2:**
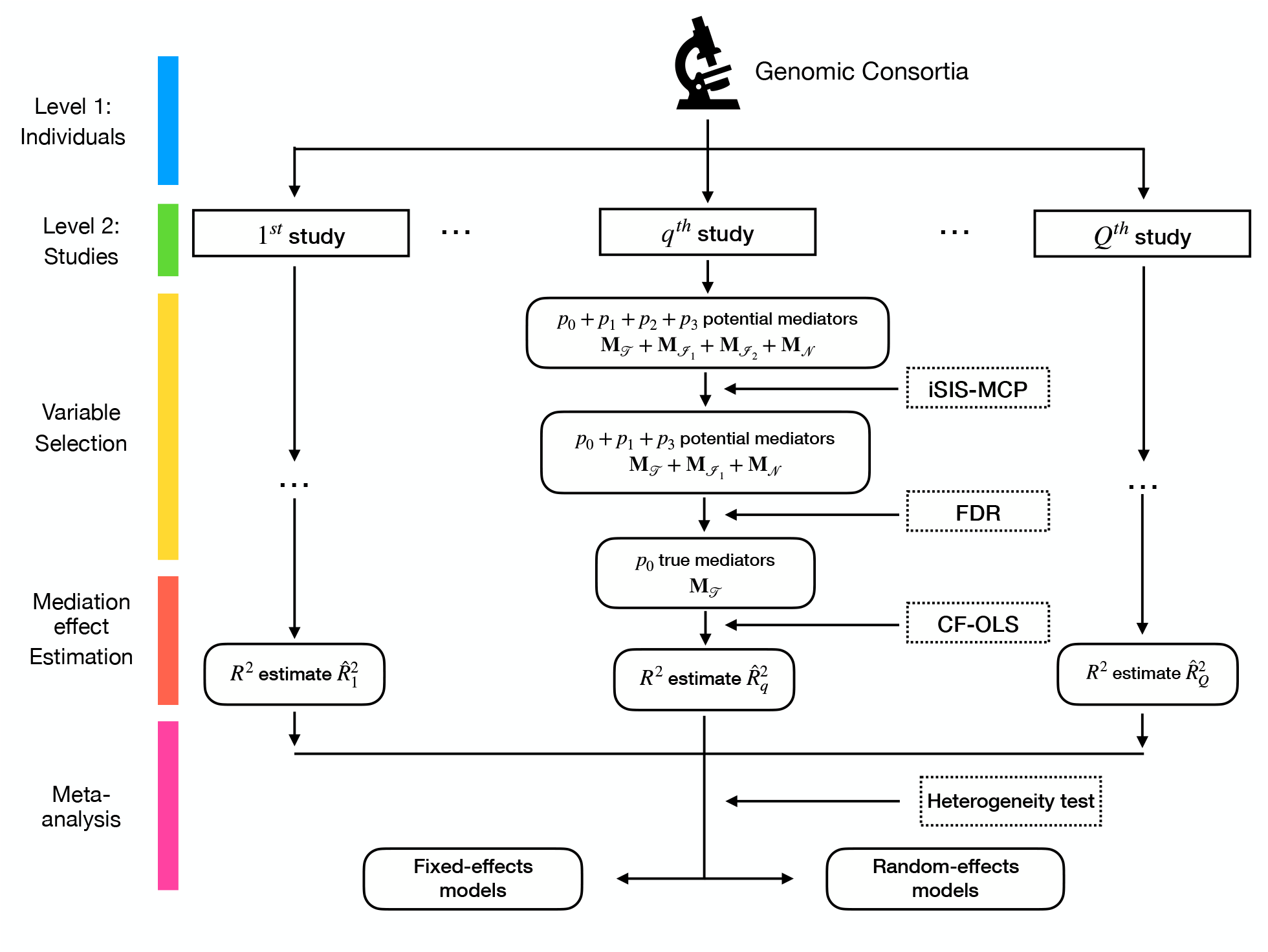
Overall workflow of meta-analysis of 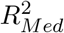 in high-dimensional mediation analysis. (*p*_0_, *p*_1_, *p*_2_, *p*_3_) refers to the number of true mediators, two types of non-mediators, and noise vari-ables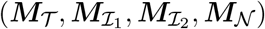, respectively.

multiple studies under high-dimensional settings. In a large-scale genomic epidemiology consortium, we first identify the potential studies relevant to our outcome of interest. Then we apply the CLOLS to each study independently, obtaining estimates of the *R*^2^-based mediation effect in each study.

Suppose that there are *Q* independent studies, each involving *n*_*q*_ participants, *q* = 1, 2, … *Q*. Let 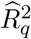 be the estimator of 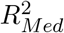 for the *q*-th study obtained using CF-OLS. Additionally, let 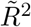 denote the estimator of 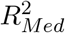 based on the individual-level data, which pools together all the studies. The

fixed-effects model in meta-analysis assumes that there is no true variability between studies beyond random sampling error. Let *R*^2^ denote the common true total mediation effect shared by all *Q* studies. The widely adopted inverse-variance estimator of 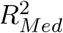 and its corresponding variance can be described as follows:

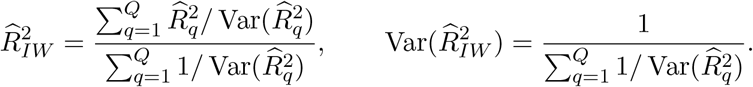

It has been shown that 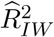 using summary statistics has the same asymptotic efficiency as using the individual-level data for all commonly used parametric models (Cox and Hinkley, 1979). Con-sequently, Var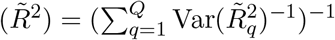, where 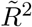 and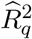, *q* = 1, 2, … *Q*, converge to *R*^2^ under

standard regularity conditions (Lin and Zeng, 2010).

The random-effects model in meta-analysis combines data from multiple studies, accounting for both within-study and between-study variability. In contrast to the fixed-effects model, the random-effects model in meta-analysis acknowledges that variations in the true effect size among studies can arise from factors beyond random sampling error. Consider the random-effects model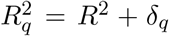, where *δ*_*q*_ *∼ N* (0, *τ* ^2^) for *q* = 1, 2, … *Q*. Let *S*_*q*_ denote the estimated variance of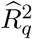. Under the assumption that the sample size *n*_*q*_ in the *q*-th study is large enough and standard regularity conditions hold, the estimate of *R*^2^ is:

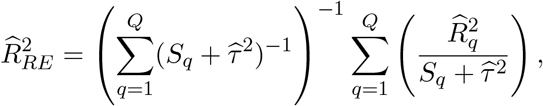

where 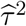 is an estimate of the between-study variance *τ* ^2^. For example, the commonly used DerSimonian and Laird estimator (DerSimonian and Laird, 1986) of *τ* ^2^ is given as

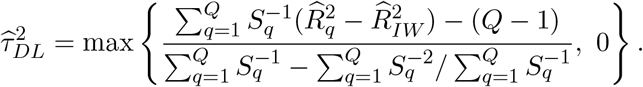

Define

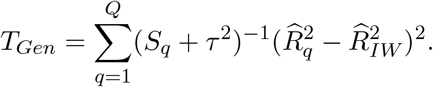

The median-unbiased Paule-Mandel estimator 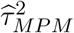 is given by the value of *τ* ^2^ such that 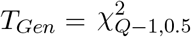 (the median of a chi-square distribution with *Q −* 1 degrees of freedom).

Following this, a heterogeneity test is conducted. The *I*^2^ statistic is a widely employed metric for quantifying heterogeneity in meta-analyses. This statistic measures the proportion of total variation in study estimates attributed to authentic between-study heterogeneity, distinct from random sampling error (Higgins and Thompson, 2002). A high *I*^2^ value (e.g., 50% to 100%) suggests high heterogeneity (Higgins et al., 2003). Based on the outcome of this assessment, we make a determination regarding the suitability of employing either the fixed-effects model or the random-effects model, guided by Cochran’s Q test (Cochran, 1954).

## 3 Simulation studies

In this section, we performed extensive simulation studies to assess the performance of the proposed meta-analysis framework for 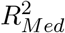 measure in high-dimensional mediation analysis. We computed coverage probability, asymptotic efficiency (i.e., standard error), bias, and empirical standard deviation of the estimator (i.e., the standard deviation of the sampling distribution of the estimator based on simulation replications). We conducted these evaluations under either fixed-effects or random-effects model, considering various high-dimensional settings to approximate real-world scenarios.

### 3.1 Simulation design

Data were simulated using the model in Eq (1), and the errors therein *ε*_1_, and *ε*_2_ independently follow the standard normal distribution. Exposure variable *X* was simulated from the standard normal distribution *N* (0, 1) and coefficient *γ* in Eq (1) was set to 3. Let (*p*_0_, *p*_1_, *p*_2_, *p*_3_) denote the number of true mediators, two types of non-mediators, and non-mediator noise variables 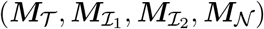, respectively.

For the fixed-effects models, iSIS was independently applied to two subsamples within each CFOLS procedure as depicted in Fig 2, for a total of 500 replications. As for the random-effects model, taking into account the sample size, we conducted 200 replications. The asymptotic standard error and bias were calculated as the means of their respective estimates across the two subsamples in the CF-OLS framework.

The performance of the two models was evaluated in various scenarios (A1)–(D1) and (A2)– (D2), respectively, each including different types or numbers of true mediators and non-mediators as follows. In scenarios A (A1 & A2), a substantial number of noise variables ***M***_*𝒩*_ were added alongside the true mediators ***M***_*𝒯*_ ; in scenarios B (B1 & B2) and scenarios C (C1 & C2), numerous non-mediators 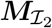 and 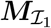 were added to the true mediators, respectively. Scenarios D (D1 & D2) examined a combination of three types of non-mediators.

The details of simulation scenarios (A)–(D) are shown as follows:

- (A) (A1)(*p*_0_, *p*_1_, *p*_2_, *p*_3_) = (150, 0, 0, 1350); (A2)(*p*_0_, *p*_1_, *p*_2_, *p*_3_) = (150, 0, 0, 4850).
- (B) (B1)(*p*_0_, *p*_1_, *p*_2_, *p*_3_) = (150, 0, 150, 1200); (B2)(*p*_0_, *p*_1_, *p*_2_, *p*_3_) = (150, 0, 150, 4700).
- (C) (C1)(*p*_0_, *p*_1_, *p*_2_, *p*_3_) = (150, 150, 0, 1200); (C2)(*p*_0_, *p*_1_, *p*_2_, *p*_3_) = (150, 150, 0, 4700).
- (D) (D1)(*p*_0_, *p*_1_, *p*_2_, *p*_3_) = (150, 150, 150, 1050); (D2)(*p*_0_, *p*_1_, *p*_2_, *p*_3_) = (150, 150, 150, 4550).

In each scenario, the same parameters ***α*** and ***β*** were simulated from a normal distribution *N* (0, 1.5^2^) across all replications, ensuring that the true 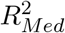 remained constant for the fixed-effects meta-analysis. In contrast, for the random-effects model, various parameters ***α*** and ***β*** were simulated from the same normal distribution *N* (0, 1.5^2^), and the true 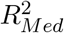 was determined as the average value among one million sets of these parameter combinations due to the true 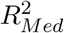 not being available in closed-form under the random-effects model. Independent variable *X* was sampled from a standard normal distribution *N* (0, 1), and coefficient *γ* in Eq (1) was set to 3. In scenarios (A1)–(D1), the independent and correlated putative mediators were considered. For independent putative mediators, the error *ξ*_*j*_ independently follows the standard normal distribution. For the putative correlated mediators, for any *i* = 1, …, *n* we consider (*ξ*_*i*1_, …, *ξ*_*ip*_)^*′*^ *∼ N* (**0**_*p×*1_, ***I***_*p*_ + **Σ**) where **Σ**_*ij*_’s are iid samples from *N* (0, 0.1^2^) for 1 ≤ *i≠ j* ≤ *p*_0_ + *p*_1_ and **Σ**_*ij*_ = 0 elsewhere. Let (*p*_0_, *p*_1_, *p*_2_, *p*_3_) denote the number of true mediators, two types of non-mediators, and noise variables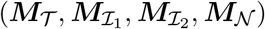, respectively. The total number of variables in ***M*** was set to *p* = 1500 for the fixed-effects model and the random-effects model in scenarios (A1)–(D1) and *p* = 5000 in scenarios (A2)–(D2). The number of studies for the fixed-effects model *Q*_*fixed*_ was selected from 1 (pooled original data) to 5, while for the random-effects model, *Q*_*random*_ was set to 5, 8, 10, 16, and 20. For the fixed-effects model, we also considered an uneven allocation of sample sizes for multiple studies (i.e., 750, 750, and 1500 for three studies), which mimics a more realistic scenario in practice. We initially generated data with *N* = 3000 and subsequently distributed them randomly across *Q*_*fixed*_ studies. In the case of the random-effects meta-analysis, we generated data with varying sample sizes for different *Q*_*random*_. We controlled the FDR level at 20% following the iSIS as shown in Fig 2.

### 3.2 Simulation results

Table 1 presents the simulation results for the fixed-effects meta-analysis of 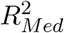 in a highdimensional setting. Overall, the fixed-effects model demonstrated good performance across all scenarios when compared to the results obtained from the original individual-level data (*Q* = 1).

**Table 1:** Simulation results using the fixed-effects model for scenarios (A1)–(D1). *N* refers to the sample size for each study. CP refers to the empirical coverage probability of 95% confidence intervals based on 200 replications. *Q*_*fixed*_ refers to the number of studies. SE refers to the average asymptotic standard error. SD refers to the empirical standard deviation of replicated estimations (ground truth).

| Scenario | $Q_{fixed}$ | $N$ | Independent mediators | | | | Correlated mediators | | | |
| --- | --- | --- | --- | --- | --- | --- | --- | --- | --- | --- |
| | | | CP<br>% | Bias<br>$\times 10^{-2}$ | SE<br>$\times 10^{-2}$ | SD<br>$\times 10^{-2}$ | CP<br>% | Bias<br>$\times 10^{-2}$ | SE<br>$\times 10^{-2}$ | SD<br>$\times 10^{-2}$ |
| A1 | 1 | 3000 | 96.5 | -0.053 | 0.455 | 0.436 | 96.5 | -0.023 | 0.413 | 0.428 |
|  | 2 | 1000 / 2000 | 96.5 | -0.019 | 0.453 | 0.440 | 92.5 | 0.002 | 0.462 | 0.495 |
|  | 2 | 1500 / 1500 | 96.5 | -0.020 | 0.453 | 0.440 | 96.0 | 0.056 | 0.511 | 0.499 |
|  | 3 | 750 / 750 / 1500 | 95.5 | 0.009 | 0.452 | 0.441 | 96.0 | 0.119 | 0.479 | 0.481 |
|  | 3 | 1000 / 1000 / 1000 | 96.5 | 0.011 | 0.452 | 0.440 | 93.5 | -0.016 | 0.388 | 0.389 |
|  | 4 | 750 / 750 / 750 / 750 | 96.0 | 0.042 | 0.451 | 0.444 | 93.0 | 0.177 | 0.496 | 0.486 |
|  | 5 | 600 / 600 / 600 / 600 / 600 | 95.0 | 0.076 | 0.450 | 0.442 | 95.5 | 0.033 | 0.487 | 0.486 |
| B1 | 1 | 3000 | 92.0 | 0.053 | 1.319 | 1.431 | 96.5 | 0.179 | 1.281 | 1.185 |
|  | 2 | 1000 / 2000 | 91.5 | 0.121 | 1.316 | 1.439 | 95.5 | 0.215 | 1.294 | 1.246 |
|  | 2 | 1500 / 1500 | 92.0 | 0.135 | 1.316 | 1.431 | 96.5 | 0.229 | 1.306 | 1.269 |
|  | 3 | 750 / 750 / 1500 | 92.0 | 0.190 | 1.313 | 1.444 | 93.5 | 0.347 | 1.316 | 1.282 |
|  | 3 | 1000 / 1000 / 1000 | 91.0 | 0.180 | 1.313 | 1.448 | 91.0 | -0.139 | 1.316 | 1.423 |
|  | 4 | 750 / 750 / 750 / 750 | 92.0 | 0.232 | 1.311 | 1.441 | 93.5 | 0.094 | 1.321 | 1.354 |
|  | 5 | 600 / 600 / 600 / 600 / 600 | 91.0 | 0.292 | 1.310 | 1.458 | 94.0 | 0.118 | 1.308 | 1.367 |
| C1 | 1 | 3000 | 94.5 | -0.060 | 0.772 | 0.785 | 91.5 | -0.111 | 0.790 | 0.851 |
|  | 2 | 1000 / 2000 | 95.0 | -0.022 | 0.770 | 0.789 | 90.5 | 0.449 | 0.744 | 0.712 |
|  | 2 | 1500 / 1500 | 94.5 | -0.020 | 0.770 | 0.782 | 93.0 | 0.061 | 0.799 | 0.836 |
|  | 3 | 750 / 750 / 1500 | 94.0 | 0.025 | 0.769 | 0.785 | 89.0 | 0.339 | 0.750 | 0.834 |
|  | 3 | 1000 / 1000 / 1000 | 94.5 | 0.023 | 0.769 | 0.795 | 92.5 | 0.327 | 0.799 | 0.775 |
|  | 4 | 750 / 750 / 750 / 750 | 94.5 | 0.071 | 0.767 | 0.789 | 95.5 | 0.249 | 0.821 | 0.788 |
|  | 5 | 600 / 600 / 600 / 600 / 600 | 94.0 | 0.105 | 0.766 | 0.797 | 86.5 | 0.479 | 0.721 | 0.748 |
| D1 | 1 | 3000 | 93.5 | -0.009 | 1.403 | 1.476 | 95.0 | 0.149 | 1.398 | 1.420 |
|  | 2 | 1000 / 2000 | 93.5 | 0.019 | 1.402 | 1.480 | 94.5 | 0.694 | 1.386 | 1.402 |
|  | 2 | 1500 / 1500 | 94.0 | 0.027 | 1.402 | 1.468 | 92.5 | 0.151 | 1.398 | 1.476 |
|  | 3 | 750 / 750 / 1500 | 93.0 | 0.046 | 1.402 | 1.498 | 93.0 | -0.552 | 1.419 | 1.599 |
|  | 3 | 1000 / 1000 / 1000 | 92.5 | 0.055 | 1.400 | 1.500 | 97.0 | -0.424 | 1.398 | 1.328 |
|  | 4 | 750 / 750 / 750 / 750 | 92.0 | 0.059 | 1.401 | 1.511 | 96.0 | 0.223 | 1.392 | 1.277 |
|  | 5 | 600 / 600 / 600 / 600 / 600 | 93.0 | 0.115 | 1.400 | 1.510 | 96.0 | 0.157 | 1.405 | 1.386 |

The empirical coverage probability of the fixed-effects model remained consistently satisfactory across all scenarios with independent mediators, closely approximating the nominal 95% level. Even in scenario (D1), where all three types of non-mediators were included and the sample sizes were down to 600 across the *Q* = 5 studies, the coverage probability remained above 90%. The coverage probability with correlated putative mediators generally performed reasonably well, maintaining above 90% except for some cases in scenario (C1). The observed bias was lowest in scenario (D1) when using the original data, but this was not the case in scenarios (A1), (B1), and (C1). In addition, the asymptotic standard errors (SEs) approximated the simulation-based standard deviations (SDs) of the estimator well, the latter of which was considered as the ground truth. A similar conclusion was reached when we included a substantial number of noise mediators ***M***_*𝒩*_ in scenarios (A2) through (D2), as detailed in the Appendix A.

Table 2 and Table 3 summarize the results based on two different between-study variance estimators in the random-effects meta-analysis of 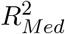 in high-dimensional settings. The coverage probability demonstrated satisfactory results when *Q*_*random*_ had a moderate value. However, for *Q*_*random*_ = 5, the coverage probability fell below 90% using the DerSimonina-Laird (DL) estimator. The empirical coverage probability of random effects models, utilizing the median-unbiased Paule-Mandel (MPM) estimator, remained consistently satisfactory across nearly all scenarios involving both independent and correlated mediators. Previous studies have indicated that achieving the correct coverage probability in random-effects meta-analysis may require *Q* = 50 studies (Zeng and Lin, 2015). Comparing the DL estimator and the MPM estimator, we observed similar bias and asymptotic standard errors. Notably, the MPM estimator tended to outperform the DL estimator in terms of coverage probability and approximation of the asymptotic SE to the simulation-based SD (ground truth), especially when the number of studies was limited (*Q*_*random*_ ≤ 10).

**Table 2:** Simulation results using the random-effects model with independent mediators for scenarios (A1)–(D1). *N* refers to the sample size for each study. CP refers to coverage probability based on 200 replications. *Q*_*random*_ refers to the number of studies. SE refers to the average asymptotic standard error. SD refers to the empirical standard deviation of replicated estimations (ground truth).

| Scenario | $Q_{random}$ | $N$ | DerSimonian-Laird | | | | Median-unbiased Paule-Mandel | | | |
| --- | --- | --- | --- | --- | --- | --- | --- | --- | --- | --- |
| | | | CP<br>% | Bias<br>$\times 10^{-2}$ | SE<br>$\times 10^{-2}$ | SD<br>$\times 10^{-2}$ | CP<br>% | Bias<br>$\times 10^{-2}$ | SE<br>$\times 10^{-2}$ | SD<br>$\times 10^{-2}$ |
| A1 | 5 | 2400 | 88.0 | 1.284 | 14.939 | 13.172 | 92.0 | 1.278 | 14.641 | 13.156 |
|  | 5 | 4000 | 87.5 | 1.245 | 14.915 | 13.134 | 92.0 | 1.242 | 14.642 | 13.124 |
|  | 8 | 1500 | 91.5 | 1.285 | 12.130 | 9.878 | 94.0 | 1.280 | 11.139 | 9.865 |
|  | 8 | 2500 | 91.0 | 1.226 | 12.082 | 9.785 | 94.0 | 1.224 | 11.148 | 9.775 |
|  | 10 | 1200 | 92.0 | 0.908 | 11.392 | 8.950 | 94.0 | 0.904 | 9.908 | 8.936 |
|  | 10 | 2000 | 92.0 | 0.849 | 11.406 | 8.971 | 94.0 | 0.849 | 9.914 | 8.958 |
|  | 16 | 750 | 93.5 | 0.482 | 9.118 | 7.748 | 94.5 | 0.483 | 7.729 | 7.741 |
|  | 16 | 1250 | 94.0 | 0.596 | 9.303 | 7.606 | 94.0 | 0.599 | 7.690 | 7.599 |
|  | 20 | 600 | 93.5 | 0.694 | 8.262 | 6.772 | 94.5 | 0.698 | 6.897 | 6.767 |
|  | 20 | 1000 | 94.0 | 0.772 | 8.509 | 6.705 | 94.0 | 0.776 | 6.862 | 6.703 |
| B1 | 5 | 2400 | 80.5 | -1.291 | 13.919 | 12.948 | 88.0 | -1.294 | 13.827 | 12.938 |
|  | 5 | 4000 | 82.0 | -1.357 | 14.000 | 12.868 | 89.0 | -1.359 | 13.858 | 12.862 |
|  | 8 | 1500 | 90.0 | -1.734 | 12.618 | 9.844 | 92.0 | -1.738 | 10.907 | 9.833 |
|  | 8 | 2500 | 89.5 | -1.752 | 12.531 | 9.860 | 91.5 | -1.755 | 10.887 | 9.852 |
|  | 10 | 1200 | 94.0 | -1.812 | 11.667 | 8.358 | 96.5 | -1.814 | 9.798 | 8.349 |
|  | 10 | 2000 | 94.0 | -1.852 | 11.636 | 8.409 | 95.0 | -1.853 | 9.786 | 8.404 |
|  | 16 | 750 | 97.0 | -1.657 | 9.496 | 6.971 | 95.5 | -1.654 | 7.680 | 6.967 |
|  | 16 | 1250 | 96.0 | -1.617 | 9.636 | 6.906 | 94.5 | -1.615 | 7.650 | 6.904 |
|  | 20 | 600 | 95.0 | -1.552 | 8.411 | 6.756 | 92.5 | -1.546 | 6.833 | 6.754 |
|  | 20 | 1000 | 94.5 | -1.511 | 8.705 | 6.712 | 92.5 | -1.507 | 6.829 | 6.713 |
| C1 | 5 | 2400 | 85.5 | 1.248 | 13.170 | 12.135 | 92.5 | 1.257 | 13.072 | 12.116 |
|  | 5 | 4000 | 84.5 | 1.276 | 13.176 | 12.055 | 93.5 | 1.282 | 13.079 | 12.043 |
|  | 8 | 1500 | 90.0 | 1.172 | 10.928 | 8.928 | 95.0 | 1.176 | 10.044 | 8.919 |
|  | 8 | 2500 | 90.0 | 1.115 | 10.726 | 8.821 | 95.5 | 1.121 | 10.043 | 8.811 |
|  | 10 | 1200 | 93.5 | 0.836 | 10.015 | 8.100 | 94.5 | 0.840 | 8.947 | 8.092 |
|  | 10 | 2000 | 93.0 | 0.849 | 9.882 | 8.094 | 95.0 | 0.854 | 8.943 | 8.084 |
|  | 16 | 750 | 94.0 | 0.473 | 8.259 | 6.934 | 95.0 | 0.478 | 6.972 | 6.928 |
|  | 16 | 1250 | 93.5 | 0.579 | 8.120 | 6.844 | 95.5 | 0.584 | 6.941 | 6.838 |
|  | 20 | 600 | 95.0 | 0.625 | 7.479 | 6.126 | 95.5 | 0.628 | 6.214 | 6.125 |
|  | 20 | 1000 | 95.0 | 0.662 | 7.381 | 6.083 | 95.5 | 0.665 | 6.189 | 6.080 |
| D1 | 5 | 2400 | 85.0 | -1.781 | 12.399 | 11.716 | 90.5 | -1.771 | 12.334 | 11.698 |
|  | 5 | 4000 | 85.5 | -1.785 | 12.427 | 11.707 | 91.0 | -1.779 | 12.329 | 11.697 |
|  | 8 | 1500 | 88.5 | -2.082 | 10.777 | 8.813 | 92.5 | -2.076 | 9.746 | 8.802 |
|  | 8 | 2500 | 89.0 | -2.123 | 10.620 | 8.853 | 93.0 | -2.117 | 9.725 | 8.845 |
|  | 10 | 1200 | 91.5 | -2.103 | 10.011 | 7.566 | 94.0 | -2.097 | 8.770 | 7.557 |
|  | 10 | 2000 | 92.0 | -2.143 | 9.823 | 7.590 | 95.0 | -2.139 | 8.741 | 7.584 |
|  | 16 | 750 | 92.5 | -1.923 | 8.101 | 6.410 | 94.0 | -1.919 | 6.857 | 6.407 |
|  | 16 | 1250 | 92.5 | -1.836 | 7.987 | 6.353 | 94.0 | -1.833 | 6.847 | 6.349 |
|  | 20 | 600 | 95.0 | -1.682 | 7.475 | 6.162 | 92.0 | -1.681 | 6.136 | 6.166 |
|  | 20 | 1000 | 95.0 | -1.689 | 7.385 | 6.072 | 91.0 | -1.687 | 6.140 | 6.074 |

**Table 3:** Simulation results using the random-effects model with correlated mediators for scenarios (A1)–(D1). *N* refers to the sample size for each study. CP refers to coverage probability based on 200 replications. *Q*_*random*_ refers to the number of studies. SE refers to the average asymptotic standard error. SD refers to the empirical standard deviation of replicated estimations (ground truth).

| Scenario | $Q_{random}$ | $N$ | DerSimonian-Laird | | | | Median-unbiased Paule-Mandel | | | |
| --- | --- | --- | --- | --- | --- | --- | --- | --- | --- | --- |
|  |  |  | CP | Bias | SE | SD | CP | Bias | SE | SD |
| | | | % | $\times 10^{-2}$ | $\times 10^{-2}$ | $\times 10^{-2}$ | % | $\times 10^{-2}$ | $\times 10^{-2}$ | $\times 10^{-2}$ |
| A1 | 5 | 2400 | 88.0 | 1.217 | 14.870 | 13.206 | 92.0 | 1.212 | 14.618 | 13.190 |
|  | 5 | 4000 | 86.0 | 1.244 | 14.911 | 13.082 | 92.5 | 1.240 | 14.612 | 13.073 |
|  | 8 | 1500 | 91.5 | 1.285 | 12.130 | 9.878 | 94.0 | 1.280 | 11.139 | 9.865 |
|  | 8 | 2500 | 91.5 | 1.229 | 12.057 | 9.829 | 94.0 | 1.226 | 11.134 | 9.821 |
|  | 10 | 1200 | 91.5 | 0.928 | 11.382 | 8.966 | 94.0 | 0.924 | 9.905 | 8.951 |
|  | 10 | 2000 | 92.0 | 0.844 | 11.409 | 8.933 | 94.5 | 0.843 | 9.944 | 8.922 |
|  | 16 | 750 | 93.5 | 0.474 | 9.252 | 7.667 | 94.5 | 0.476 | 7.743 | 7.658 |
|  | 16 | 1250 | 92.5 | 0.594 | 9.316 | 7.579 | 93.5 | 0.595 | 7.691 | 7.574 |
|  | 20 | 600 | 93.5 | 0.614 | 8.366 | 6.764 | 95.0 | 0.619 | 6.911 | 6.759 |
|  | 20 | 1000 | 92.5 | 0.722 | 8.556 | 6.712 | 95.5 | 0.726 | 6.865 | 6.709 |
| B1 | 5 | 2400 | 81.5 | -1.292 | 13.977 | 12.917 | 88.5 | -1.295 | 13.857 | 12.908 |
|  | 5 | 4000 | 81.5 | -1.393 | 13.949 | 12.811 | 89.0 | -1.394 | 13.900 | 12.804 |
|  | 8 | 1500 | 90.0 | -1.734 | 12.618 | 9.844 | 92.0 | -1.738 | 10.907 | 9.833 |
|  | 8 | 2500 | 89.5 | -1.756 | 12.582 | 9.841 | 91.5 | -1.758 | 10.918 | 9.833 |
|  | 10 | 1200 | 94.0 | -1.859 | 11.720 | 8.322 | 96.0 | -1.861 | 9.813 | 8.314 |
|  | 10 | 2000 | 93.0 | -1.949 | 11.657 | 8.338 | 94.5 | -1.950 | 9.783 | 8.333 |
|  | 16 | 750 | 96.0 | -1.800 | 9.558 | 6.949 | 95.0 | -1.798 | 7.694 | 6.945 |
|  | 16 | 1250 | 96.5 | -1.686 | 9.650 | 6.814 | 95.5 | -1.684 | 7.656 | 6.812 |
|  | 20 | 600 | 95.0 | -1.677 | 8.543 | 6.785 | 92.5 | -1.671 | 6.857 | 6.784 |
|  | 20 | 1000 | 93.5 | -1.642 | 8.738 | 6.780 | 93.0 | -1.638 | 6.853 | 6.780 |
| C1 | 5 | 2400 | 85.5 | 1.272 | 13.170 | 12.121 | 92.0 | 1.281 | 13.089 | 12.101 |
|  | 5 | 4000 | 86.0 | 1.099 | 13.143 | 12.047 | 92.5 | 1.112 | 13.089 | 12.023 |
|  | 8 | 1500 | 89.5 | 1.092 | 10.881 | 8.935 | 94.5 | 1.099 | 10.062 | 8.921 |
|  | 8 | 2500 | 89.0 | 1.166 | 10.687 | 8.917 | 95.0 | 1.176 | 10.037 | 8.901 |
|  | 10 | 1200 | 93.0 | 0.753 | 10.014 | 8.107 | 94.5 | 0.759 | 8.964 | 8.096 |
|  | 10 | 2000 | 91.5 | 0.786 | 9.762 | 8.059 | 94.5 | 0.792 | 8.930 | 8.050 |
|  | 16 | 750 | 93.0 | 0.435 | 8.221 | 6.909 | 95.5 | 0.440 | 6.987 | 6.903 |
|  | 16 | 1250 | 91.5 | 0.465 | 7.879 | 6.946 | 95.0 | 0.474 | 6.977 | 6.933 |
|  | 20 | 600 | 95.0 | 0.495 | 7.506 | 6.149 | 95.0 | 0.499 | 6.227 | 6.147 |
|  | 20 | 1000 | 92.0 | 0.512 | 7.220 | 6.172 | 95.0 | 0.518 | 6.221 | 6.165 |
| D1 | 5 | 2400 | 85.0 | -1.772 | 12.416 | 11.691 | 90.0 | -1.762 | 12.344 | 11.674 |
|  | 5 | 4000 | 85.5 | -1.913 | 12.438 | 11.680 | 90.0 | -1.907 | 12.332 | 11.670 |
|  | 8 | 1500 | 88.5 | -2.165 | 10.750 | 8.891 | 91.5 | -2.156 | 9.753 | 8.878 |
|  | 8 | 2500 | 87.5 | -2.152 | 10.596 | 8.856 | 94.0 | -2.147 | 9.726 | 8.848 |
|  | 10 | 1200 | 91.5 | -2.168 | 10.004 | 7.559 | 94.0 | -2.160 | 8.792 | 7.549 |
|  | 10 | 2000 | 91.0 | -2.065 | 9.765 | 7.666 | 94.5 | -2.056 | 8.752 | 7.651 |
|  | 16 | 750 | 93.0 | -1.962 | 8.105 | 6.425 | 93.5 | -1.958 | 6.878 | 6.421 |
|  | 16 | 1250 | 91.0 | -2.007 | 7.965 | 6.465 | 94.5 | -2.002 | 6.894 | 6.460 |
|  | 20 | 600 | 94.5 | -1.797 | 7.483 | 6.168 | 92.0 | -1.795 | 6.157 | 6.171 |
|  | 20 | 1000 | 94.0 | -1.769 | 7.273 | 6.119 | 91.5 | -1.766 | 6.166 | 6.118 |

## 4 Real Data Applications

In this section, we describe the cohorts in the TOPMed program, including subject recruitment, ethnic diversity, and high-throughput technologies. We then apply our proposed meta-analysis framework to cardiovascular disease (CVD) traits in these studies as a proof of concept.

### 4.1 Heterogeneity in cohorts, ethnic representations, and mRNA profiling technologies

The Trans-Omics for Precision Medicine (TOPMed) program represents a groundbreaking initiative launched by the National Heart, Lung, and Blood Institute (NHLBI) with the vision of aggregating whole-genome sequencing (WGS) and other omics data from more than 85 population studies (Taliun et al., 2021; Hu et al., 2021). The program’s objective is to uncover the genetic and molecular foundations associated with heart, lung, blood, and sleep disorders (Hu et al., 2021). The Framingham Heart Study (FHS) began the recruitment for the Offspring cohort in 1971, which comprises the children of the Original cohort and their spouses. The Offspring cohort consists of 5,124 individuals, of which 52% are female(Kannel et al., 1979). In 2002, FHS initiated the Third-Generation cohort, encompassing the children of the Offspring cohort, consisting of 4,095 participants of which 54% are female (Mahmood et al., 2014). The vast majority of the FHS participants are Non-Hispanic Whites. Another active and comprehensive cohort study from the TOPMed is the Multi-Ethnic Study of Atherosclerosis (MESA), encompassing 6,814 individuals aged 45 to 84 from six U.S. communities (Olson et al., 2016). MESA is dedicated to unraveling the risk factors that contribute to the development of CVD, particularly focusing on atherosclerosis, across 4 ethnic groups, including Non-Hispanic Whites, African Americans, Hispanics, and Chinese Americans (Lakoski et al., 2006).

The transcriptome encompasses all messenger RNAs (mRNAs)/transcripts in a cell during a specific stage or condition. It is crucial for deciphering the genome’s functional elements, understanding cellular components, and gaining insights into development and disease (Clark et al., 2002). Typically, hybridization-based microarray gene expression profiling is cost-effective and high-throughput but depends on current genomic knowledge (Sud et al., 2017). Conversely, RNA sequencing (RNA-seq) facilitates the identification of new gene transcripts and non-coding RNAs (Wang et al., 2009). Thanks to the decreasing cost of next-generation sequencing technologies, RNAseq has become more affordable and feasible in large-scale studies such as the FHS and MESA.

### 4.2 Applications to CVD Traits

Hypertension stands as the primary contributor to global CVD and premature mortality (Mills et al., 2020). In 2010, 31.1% of the global adult population (1.39 billion), were diagnosed with hypertension, characterized by a systolic blood pressure (BP) of ≥140 mmHg and/or a diastolic BP of ≥90 mmHg (Mills et al., 2016). Parallel to the observed increase in hypertension prevalence, the estimated counts of all-cause and CVD mortalities associated with high BP showed a significant rise from 1990 to 2015 (Forouzanfar et al., 2017). On the other hand, previous epidemiological studies consistently identified inverse linear associations between high-density lipoprotein cholesterol (HDLC) levels and the risks associated with CVD and mortality (Gordon et al., 1989; Wilson et al., 1988; Castelli et al., 1986). Meanwhile, many findings have highlighted notable sexual dimorphism in HDL-C levels and functionality (Palmisano et al., 2018; Wang et al., 2011).

We applied our developed meta-analysis method to the FHS Offspring cohort, the FHS ThirdGeneration cohort, and the MESA cohorts to estimate the mediation effects of gene expression on age-related variation in systolic BP and sex-related variation in HDL-C. Systolic BP was determined by averaging two physician-taken readings (rounded to the nearest 2 mm Hg). For individuals on anti-hypertensive medication, an adjustment was made by adding 15 mm Hg to their reading (Tobin et al., 2005). HDL-C was measured from EDTA plasma (in mg/dL), and age was recorded based on the participant’s age at the time of examination. The covariates included body mass index (BMI, expressed in *kg/m*^2^), dichotomized smoking status (current smoker or non-smoker), and dichotomized drinking status (never or ever). When a variable, either age or sex, was considered the exposure of interest, the other was incorporated as a covariate in the model by regressing out the covariate and working on the residuals subsequently (Xu et al., 2023). In the FHS Offspring and Third-Generation cohorts, expression profiling for 17,873 genes/transcripts was conducted using the Affymetrix Human Exon 1.0 ST GeneChip, derived from whole blood mRNA (Joehanes et al., 2012). On the other hand, as part of the TOPMed program, RNA-seq was performed on whole blood in these FHS cohorts using the Illumina NovaSeq system profiling expression of over 40,000 transcripts (Keshawarz et al., 2023). In the meta analysis, we only included nonoverlapping participants between the microarray and RNAseq platforms. For MESA’s four ethnicity cohorts, RNA-seq measured expression profiles for more than 30,000 transcripts (Hu et al., 2023). Exposures, covariates, and gene expression levels were extracted from the FHS Offspring cohort’s 8^*th*^ examination, the FHS Third-Generation cohort’s 2^*nd*^ examination, and the MESA cohort’s 1^*st*^ examination. Phenotype data was gathered from the Offspring cohort’s 9^*th*^ examination, the ThirdGeneration cohort’s 3^*rd*^ examination, and the MESA cohort’s 1^*st*^ examination to ensure temporal order that the exposure affects the mediators which in turn precedes the outcome (Kraemer et al., 2002).

In Fig 3A, we present fixed-effects meta-analysis results investigating the total mediation effect of gene expression in the relationship between age and systolic BP across 8 cohorts from the TOPMed program. Both the sample size and the number of profiled transcripts varied across cohorts. We employed the CF-OLS on each cohort to identify the true mediators, subsequently obtaining the 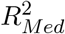 estimate along with its 95% confidence interval. We also considered the Shared Over Simple (SOS) measure. Defined as 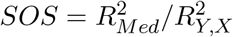, this measure represents the standardized variance in the outcome related to the exposure that intersects with the mediators (Lindenberger and Pötter, 1998). For example, in the FHS Third-Generation RNA-seq cohort, a total of 1635 subjects with complete data were included in the analysis. We applied the CF-OLS procedure to perform variable selection out of the 47437 transcripts measured using RNA-seq, in which 117 and 109 transcripts remained in each of the two subsamples for the estimation. We then estimated that 3.5% (95% CI = (1.5%, 5.5%)) of systolic BP variation was attributable to the indirect effect of age, mediated by gene expression, i.e., SOS = 29.9% (95% CI = (15.9%, 43.8%)) of the age-related variation in systolic BP was mediated by gene expression in the FHS Third-Generation RNA-seq cohort. Furthermore, we computed the *I*^2^ statistic to quantify the extent of heterogeneity, offering a measure of the degree of inconsistency in results across cohorts (Higgins et al., 2003). Given the lack of heterogeneity (*I*^2^ = 0%, *p* = 0.76), we opted for the proposed fixed-effects model to combine the total mediation effects from diverse cohorts. Consequently, 3.6% (95% CI = (2.6%, 4.6%)) of the variance in systolic BP was explained by age through gene expression (SOS = 30.2% (95% CI = (23.4%, 37.1%))). Fixed-effects meta-analysis yielded comparable point estimation and confidence intervals as the previous study, suggesting that the new method effectively gives and combines reliable estimates across diverse cohorts (Xu et al., 2023).

**Figure 3:**
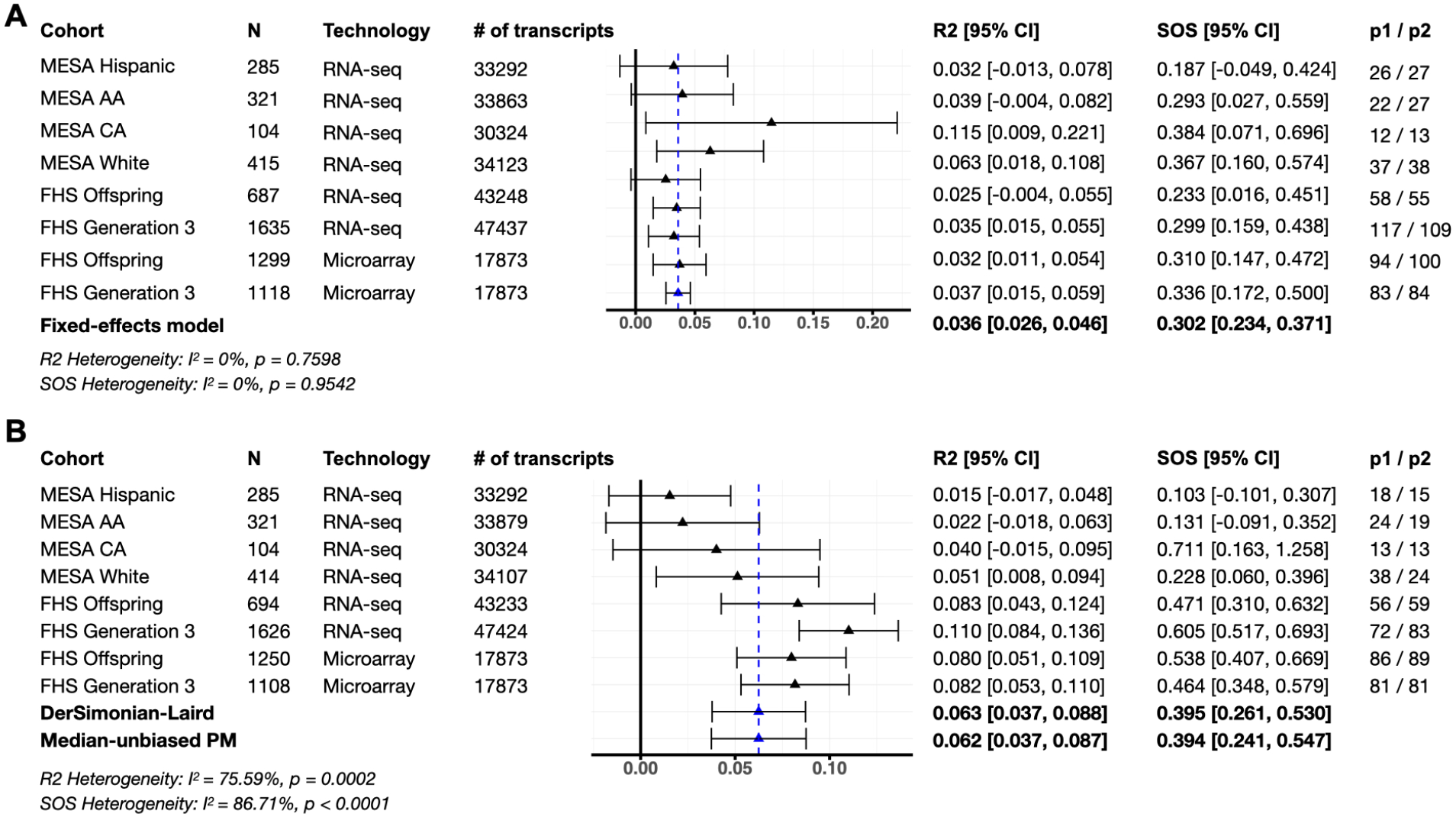
Meta-analysis results using the CF-OLS in 8 different cohorts from the NHLBI TOPMed program. (A) Fixed-effects model results of mediation effect of gene expression between age and systolic BP and (B) random-effects model results of mediation effect of gene expression between sex and HDL-C. N refers to the sample size. Technology refers to the high-throughput gene expression profiling technology. # of transcripts refers to the number of genes measured from the gene expression profiling. p1 */* p2 refers to the number of transcripts selected in the first and second subsample, respectively. R2 refers to the total mediation effect 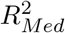. CI refers to the confidence interval. CA refers to the Chinese American. AA refers to African American.

Fig 3B displays the results of mediation analysis of gene expression in the relationship between sex and HDL-C, including the same 8 cohorts from the TOPMed program. With the observed heterogeneity between cohorts (*I*^2^ = 75.59%, *p* < 0.01), we chose the random-effects meta-analysis for 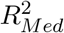. For example, the FHS Third-Generation RNA-seq cohort exhibited a total mediation effect of 11.0% (95% CI = (8.4%, 13.6%)), which is nearly tenfold greater than the 1.5% (95% CI = (-1.7%, 4.8%)) observed in the MESA Hispanic cohort. Using the proposed random-effects metaanalysis model, we estimated that 6.3% (95% CI = (3.7%, 8.7%)) of HDL-C variation using the DL estimator could be explained by sex through the mediation of gene expression (SOS = 39.4% (95% CI = (24.1%, 53.0%))). The MPM estimator, shown to have an edge over the DL estimator when the number of studies was limited (Table 3), was employed to estimate the between-study variance (Sidik and Jonkman, 2005). The results were comparable.

Since the sample sizes in four MESA cohorts were less than 500, we combined them into a single cohort (*N* = 1125 for systolic BP outcome and *N* = 1124 for HDL-C outcome) and then conducted the analysis, as shown in Appendix B. Given the observed lack of heterogeneity between cohorts for the systolic BP outcome (*I*^2^ = 0%, *p* = 0.6851), the fixed-effects model similarly concluded that 3.6% (95% CI = (2.6%, 4.7%)) of the variance in systolic BP could be explained by age through gene expression. For HDL-C, the DL estimator indicated that 8.1% (95% CI = (5.9%, 10.3%)) of the variation could be explained by sex through gene expression, with the MPM estimator providing a nearly identical estimate of 8.1% (95% CI = (6.0%, 10.2%)).

To investigate the biological pathways involved, we conducted a pathway enrichment analysis on the mediator genes selected in each cohort. We then carried out a meta-analysis of the enrichment pvalues for the pathways (see Appendix B). This analysis utilized the sample size-weighted Stouffer’s combination of p-values (Willer et al., 2010). We observed that there were more enriched pathways (meta-analysis p-value ≤ 0.05) for HDL-C than for SBP. Notably, the endocytosis pathway plays a critical role in cellular processes and could influence lipid metabolism, including HDL-C levels (Fruhwürth et al., 2013).

## 5 Discussion

We have introduced a novel and efficient method for conducting fixed-effects and random-effects meta-analyses for the 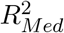 total mediation effect in high-dimensional settings. This method only requires summary statistics and accounts for between-study heterogeneity. Our approach incorporates iSIS-MCP into two subsamples to eliminate the non-mediators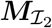. We then apply FDR control to filter out the non-mediators 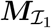 and noise variable ***M***_*𝒩*_ . We then obtain the point estimate and asymptotic standard error of 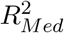 via the CF-OLS procedure. Depending on the results of the heterogeneity test, we subsequently perform either fixed-effects or random-effects meta-analysis.

Based on our simulations, we demonstrate that the relative efficiency and coverage probability achieved using summary statistics are comparable with those obtained from the original individual-level data in a fixed-effects meta-analysis. Additionally, our simulations indicate that conducting a meta-analysis for total mediation effect is reliable with a minimum sample size of around 300 in each study. This is particularly applicable when the study sizes are comparable, or when there are larger studies that can offset those with more limited sample sizes. Furthermore, in the more realistic scenario where the assumption of a common effect size across all studies no longer holds, the random-effects model maintains an acceptable coverage probability when the number of studies is relatively large, for example, larger than 10, which holds for most large-scale genomic consortia, such as the TOPMed program with over 85 studies and The Global Lipids Genetics Consortium with over 200 studies (Graham et al., 2021).

In the TOPMed program, as a proof of concept we applied our proposed new meta-analysis framework across various FHS and MESA cohorts to assess the mediation effects of gene expression on age-related variation in systolic BP and sex-related variation in HDL-C. Our findings closely align with results derived from the original individual-level data with much less computational cost, highlighting the efficiency of our method in handling the computational burden caused by large-scale studies. This is particularly applicable to mediation analysis in large-scale biobanks, such as the UK Biobank of over a half million participants with diverse ethnicity and multi-omics profiling based on different platforms. Meta-analysis can be an appealing alternative to analysis of the entire dataset at once in terms of computational feasibility (Li et al., 2023) and evaluating potential heterogeneity across risk factors for common diseases and genomic profiling technologies, as demonstrated in our application to the FHS and MESA cohorts.

## Availability and Implementation

The proposed meta-analysis method has been implemented in R package `MetaR2M` available on `GitHub` at https://github.com/zhichaoxu04/MetaR2M and will be submitted to R/CRAN. The data sets used for the analyses described in this manuscript were obtained from NIH/dbGaP at https://www.ncbi.nlm.nih.gov/gap/ through accession numbers phs000007, phs000974, phs000209, and phs001416.

## Acknowledgements

This research was partially supported by National Institutes of Health (NIH) grant R01HL116720. The Framinham Heart Study (FHS) is conducted and supported by the National Heart, Lung, and Blood Institute (NHLBI) in collaboration with Boston University. The Multi-Ethnic Study of Atherosclerosis (MESA) is conducted and supported by the NHLBI in collaboration with MESA investigators. This manuscript was not prepared in collaboration with investigators in the FHS or MESA and does not necessarily reflect the opinions or views of the FHS, Boston University, the MESA, or the NHLBI.

## Appendix A Additional simulations and details

Table 4 presents the simulation results for the fixed-effects meta-analysis of 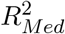 in a highdimensional setting in scenarios (A2)–(D2). Overall, the fixed-effects model demonstrated good performance across all scenarios when compared to the results obtained from the original individual-level data (*Q* = 1).

**Table 4:** Simulation results using the fixed-effects model with independent mediators for scenarios (A2)–(D2). CP refers to the empirical coverage probability of 95% confidence intervals based on 200 replications. *Q*_*fixed*_ refers to the number of studies. SE refers to the average asymptotic standard error. SD refers to the empirical standard deviation of replicated estimations.

| Scenario | $Q_{fixed}$ | $N$ | CP<br>% | Bias<br>$\times 10^{-2}$ | SE<br>$\times 10^{-2}$ | SD<br>$\times 10^{-2}$ |
| --- | --- | --- | --- | --- | --- | --- |
| A2 | 1 | 3000 | 94.5 | 0.006 | 0.452 | 0.480 |
|  | 2 | 1000 / 2000 | 94.0 | 0.044 | 0.450 | 0.483 |
|  | 2 | 1500 / 1500 | 93.0 | 0.044 | 0.450 | 0.482 |
|  | 3 | 750 / 750 / 1500 | 93.0 | 0.076 | 0.449 | 0.481 |
|  | 3 | 1000 / 1000 / 1000 | 93.5 | 0.070 | 0.449 | 0.480 |
|  | 4 | 750 / 750 / 750 / 750 | 91.0 | 0.103 | 0.448 | 0.484 |
|  | 5 | 600 / 600 / 600 / 600 / 600 | 91.0 | 0.139 | 0.447 | 0.488 |
| B2 | 1 | 3000 | 97.5 | 0.159 | 1.316 | 1.185 |
|  | 2 | 1000 / 2000 | 97.5 | 0.229 | 1.314 | 1.176 |
|  | 2 | 1500 / 1500 | 97.0 | 0.218 | 1.314 | 1.195 |
|  | 3 | 750 / 750 / 1500 | 97.5 | 0.281 | 1.311 | 1.200 |
|  | 3 | 1000 / 1000 / 1000 | 97.5 | 0.285 | 1.311 | 1.187 |
|  | 4 | 750 / 750 / 750 / 750 | 97.5 | 0.317 | 1.309 | 1.205 |
|  | 5 | 600 / 600 / 600 / 600 / 600 | 97.5 | 0.362 | 1.308 | 1.199 |
| C2 | 1 | 3000 | 93.5 | 0.058 | 0.768 | 0.768 |
|  | 2 | 1000 / 2000 | 93.0 | 0.107 | 0.766 | 0.774 |
|  | 2 | 1500 / 1500 | 94.0 | 0.103 | 0.766 | 0.766 |
|  | 3 | 750 / 750 / 1500 | 93.5 | 0.151 | 0.764 | 0.768 |
|  | 3 | 1000 / 1000 / 1000 | 93.0 | 0.149 | 0.765 | 0.768 |
|  | 4 | 750 / 750 / 750 / 750 | 92.5 | 0.198 | 0.763 | 0.777 |
|  | 5 | 600 / 600 / 600 / 600 / 600 | 92.5 | 0.249 | 0.761 | 0.771 |
| D2 | 1 | 3000 | 94.0 | 0.152 | 1.405 | 1.395 |
|  | 2 | 1000 / 2000 | 93.5 | 0.175 | 1.403 | 1.397 |
|  | 2 | 1500 / 1500 | 93.5 | 0.175 | 1.403 | 1.394 |
|  | 3 | 750 / 750 / 1500 | 93.5 | 0.198 | 1.403 | 1.398 |
|  | 3 | 1000 / 1000 / 1000 | 94.0 | 0.199 | 1.402 | 1.413 |
|  | 4 | 750 / 750 / 750 / 750 | 94.0 | 0.213 | 1.402 | 1.414 |
|  | 5 | 600 / 600 / 600 / 600 / 600 | 93.0 | 0.231 | 1.400 | 1.426 |

The details of simulation scenarios (A2)–(D2) are shown as follows:

- (A2)(*p*_0_, *p*_1_, *p*_2_, *p*_3_) = (150, 0, 0, 4850).
- (B2)(*p*_0_, *p*_1_, *p*_2_, *p*_3_) = (150, 0, 150, 4700).
- (C2)(*p*_0_, *p*_1_, *p*_2_, *p*_3_) = (150, 150, 0, 4700).
- (D2)(*p*_0_, *p*_1_, *p*_2_, *p*_3_) = (150, 150, 150, 4550).

## Appendix B Details and supplement of the applications

We applied the FDR-adjusted p-value to filter out selected genes not associated with the exposures, setting the FDR cutoff point at 0.2. We conducted pathway enrichment analysis using the Database for Annotation, Visualization and Integrated Discovery (DAVID) (Dennis Jr et al., 2003) to evaluate the significance of those mediating genes enriched in specific pathways.

Table 5 presents the meta-analysis results by sample size-weighted Stouffer’s combination of pvalues (Willer et al., 2010) for pathways identified for systolic BP from the Kyoto Encyclopedia of Genes and Genomes (KEGG) (Kanehisa and Goto, 2000), which were ranked by the meta-analysis p-value. Table 6 lists the pathways identified for HDL-C.

**Table 5:** The Top 10 significant pathways and p-value identified for systolic blood pressure. Micro GEN3 refers to the FHS Third Generation cohort with microarray gene expression profiling. Micro OFF refers to the FHS Offspring cohort with microarray gene expression profiling. RNA GEN3 refers to the FHS Third Generation cohort with RNAseq gene expression profiling. RNA OFF refers to the FHS Offspring cohort with RNAseq gene expression profiling. Overall refers to the overall p-value using the sample size based METAL.

| Pathway | Count | Genes | Micro_GEN3 | Micro_OFF | RNA_GEN3 | RNA_OFF | MESA | Overall |
| --- | --- | --- | --- | --- | --- | --- | --- | --- |
| hsa04810:Regulation of actin cytoskeleton | 3 | CXCL12, NCKAP1L, ITGAL | 0.536 | 0.758 | 0.145 | 1.000 | 0.025 | 0.030 |
| hsa03013:Nucleocytoplasmic transport | 3 | NUP214, RNPS1, IPO5 | 0.181 | 0.328 | 1.000 | 1.000 | 1.000 | 0.296 |
| hsa04814:Motor proteins | 3 | KIF1C, KIF17, ACTG1 | 0.754 | 0.624 | 0.274 | 1.000 | 1.000 | 0.344 |
| hsa01100:Metabolic pathways | 5 | UGT8, RRM2, ATIC, HYAL4, PGM2L1 | 0.998 | 0.626 | 0.932 | 0.858 | 0.335 | 0.448 |
| hsa04141:Protein processing in endoplasmic reticulum | 5 | GANAB, RPN1, DERL1, STT3A, SEC24C | 1.000 | 0.113 | 1.000 | 1.000 | 1.000 | 0.456 |
| hsa05132:Salmonella infection | 6 | TUBB6, ARHGEF26, NCKAP1L, VPS39, S100A10, DNM2 | 1.000 | 0.715 | 0.709 | 1.000 | 0.456 | 0.487 |
| hsa04120:Ubiquitin mediated proteolysis | 2 | HUWE1, BIRC6 | 0.642 | 0.790 | 0.533 | 1.000 | 1.000 | 0.511 |
| hsa05203:Viral carcinogenesis | 4 | RASA2, UBR4, CHD4, CCR4 | 0.773 | 0.382 | 1.000 | 1.000 | 1.000 | 0.591 |
| hsa04380:Osteoclast differentiation | 3 | FCGR3A, ATP2A2, FYN | 1.000 | 0.432 | 1.000 | 1.000 | 1.000 | 0.712 |
| hsa05200:Pathways in cancer | 6 | LAMA5, CXCL12, LAMA1, LAMC3, LAMA3, COL4A6 | 0.901 | 0.691 | 0.946 | 1.000 | 0.879 | 0.731 |

**Table 6:** The Top 10 significant pathways and p-value identified for HDL-C. Micro GEN3 refers to the FHS Third Generation cohort with microarray gene expression profiling. Micro OFF refers to the FHS Offspring cohort with microarray gene expression profiling. RNA GEN3 refers to the FHS Third Generation cohort with RNAseq gene expression profiling. RNA OFF refers to the FHS Offspring cohort with RNAseq gene expression profiling. Overall refers to the overall p-value using the sample size based METAL.

| Pathway | Count | Genes | Micro_GEN3 | Micro_OFF | RNA_GEN3 | RNA_OFF | MESA | Overall |
| --- | --- | --- | --- | --- | --- | --- | --- | --- |
| hsa04144:Endocytosis | 6 | GRK2, GRK6, VPS26B, TGFBR2, DNM2, HLA-E | 0.262 | 0.556 | 0.113 | 0.147 | 0.311 | 0.011 |
| hsa04072:Phospholipase D signaling pathway | 2 | DNM2, PIK3R5 | 0.457 | 0.669 | 0.096 | 0.008 | 1.000 | 0.020 |
| hsa04010:MAPK signaling pathway | 5 | MAP3K2, CACNG8, JUN, RAS-GRF2, NF1 | 0.203 | 0.182 | 0.658 | 0.214 | 1.000 | 0.066 |
| hsa04810:Regulation of actin cytoskeleton | 3 | CXCL12, NCKAP1L, ITGAL | 0.693 | 0.821 | 0.821 | 0.007 | 0.276 | 0.070 |
| hsa04910:Insulin signaling pathway | 2 | SREBF1, EXOC7 | 0.764 | 0.269 | 0.080 | 0.521 | 1.000 | 0.073 |
| hsa05012:Parkinson disease | 3 | PSMC5, GPR37, SLC25A31 | 0.941 | 0.046 | 0.865 | 0.052 | 1.000 | 0.085 |
| hsa04151:PI3K-Akt signaling pathway | 6 | LAMA5, FLT1, LAMA1, LAMC3, LAMA3, COL4A6 | 0.317 | 0.750 | 0.125 | 0.574 | 0.809 | 0.090 |
| hsa01100:Metabolic pathways | 6 | PIGT, GPAT4, ME1, UQCR11, GAPDH, ATP6V0C | 0.756 | 0.351 | 0.830 | 0.055 | 0.559 | 0.109 |
| hsa04630:JAK-STAT signaling pathway | 2 | IL21R, IL5RA | 0.518 | 0.711 | 0.034 | 1.000 | 1.000 | 0.115 |
| hsa04150:mTOR signaling pathway | 2 | STK11, FZD6 | 1.000 | 0.689 | 0.028 | 0.568 | 1.000 | 0.122 |

Fig 4 and Fig 5 present the application results using 5 cohorts for the systolic BP and HDL-C. Given the observed lack of heterogeneity between cohorts for the systolic BP outcome (*I*^2^ = 0%, *p* = 0.6851), the fixed-effects model similarly concluded that 3.6% (95% CI = (2.6%, 4.7%)) of the variance in systolic BP could be explained by age through gene expression. For HDL-C, the observed heterogeneity between cohorts for the HDL-C outcome (*I*^2^ = 57.02%, *p* = 0.0539) is moderate even though it is not statistically significant. The DL estimator indicated that 8.1% (95% CI = (5.9%, 10.3%)) of the variation could be explained by sex through gene expression, with the MPM estimator providing a nearly identical estimate of 8.1% (95% CI = (6.0%, 10.2%)). We also performed the fixed-effects model and concluded that 8.8% (95% CI = (7.5%, 10.1%)) of the variation could be explained by sex through gene expression.

**Figure 4:**
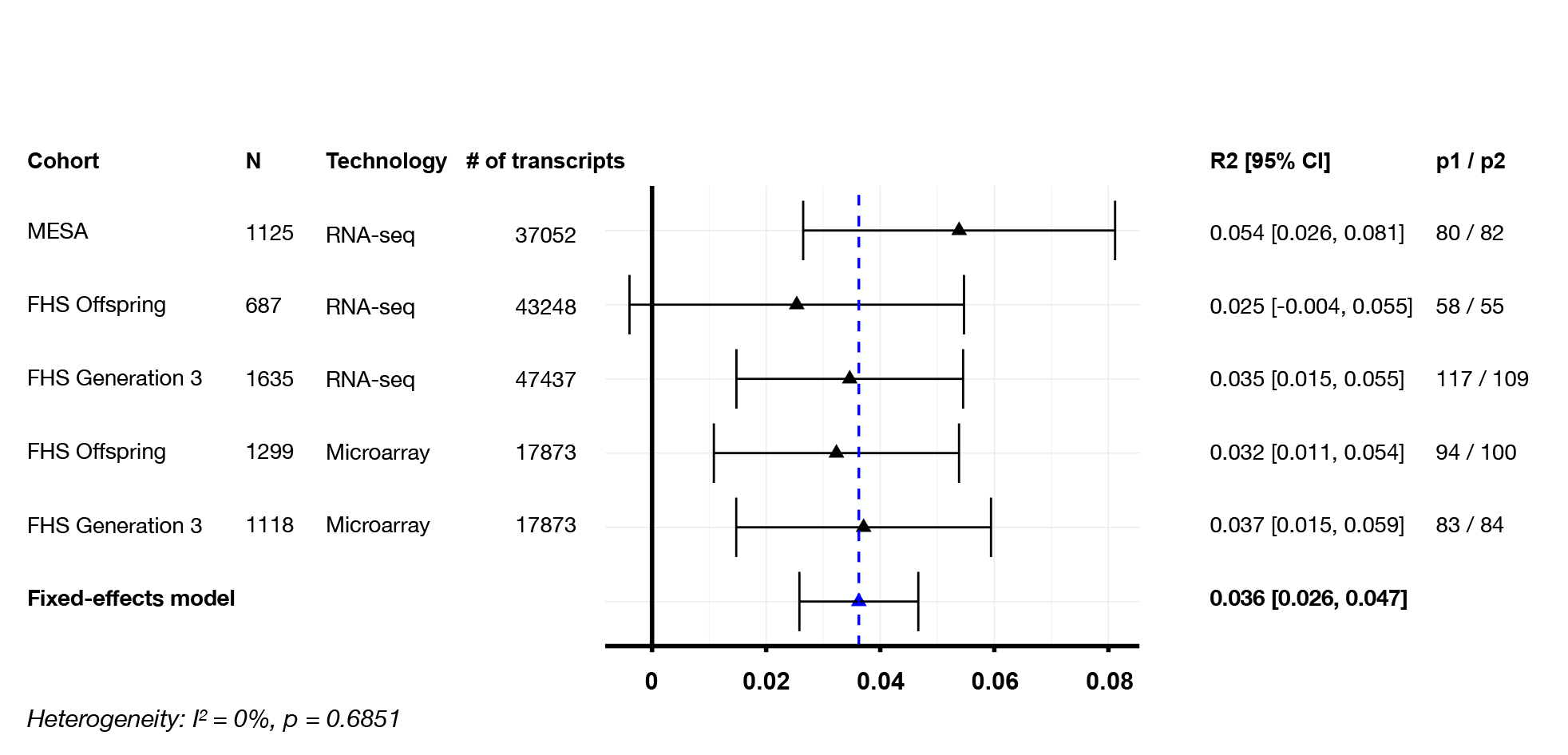
Fixed-effects meta-analysis results of mediation effect of gene expression between age and systolic BP using the CF-OLS in 5 different cohorts from the NHLBI TOPMed program. *N* refers to the sample size. # of transcripts refers to the number of genes measured from the gene expression profiling. *p*1*/p*2 refers to the number of transcripts selected in the first and second subsamples, respectively. R2 refers to the total mediation effect 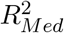. CI refers to the confidence interval.

**Figure 5:**
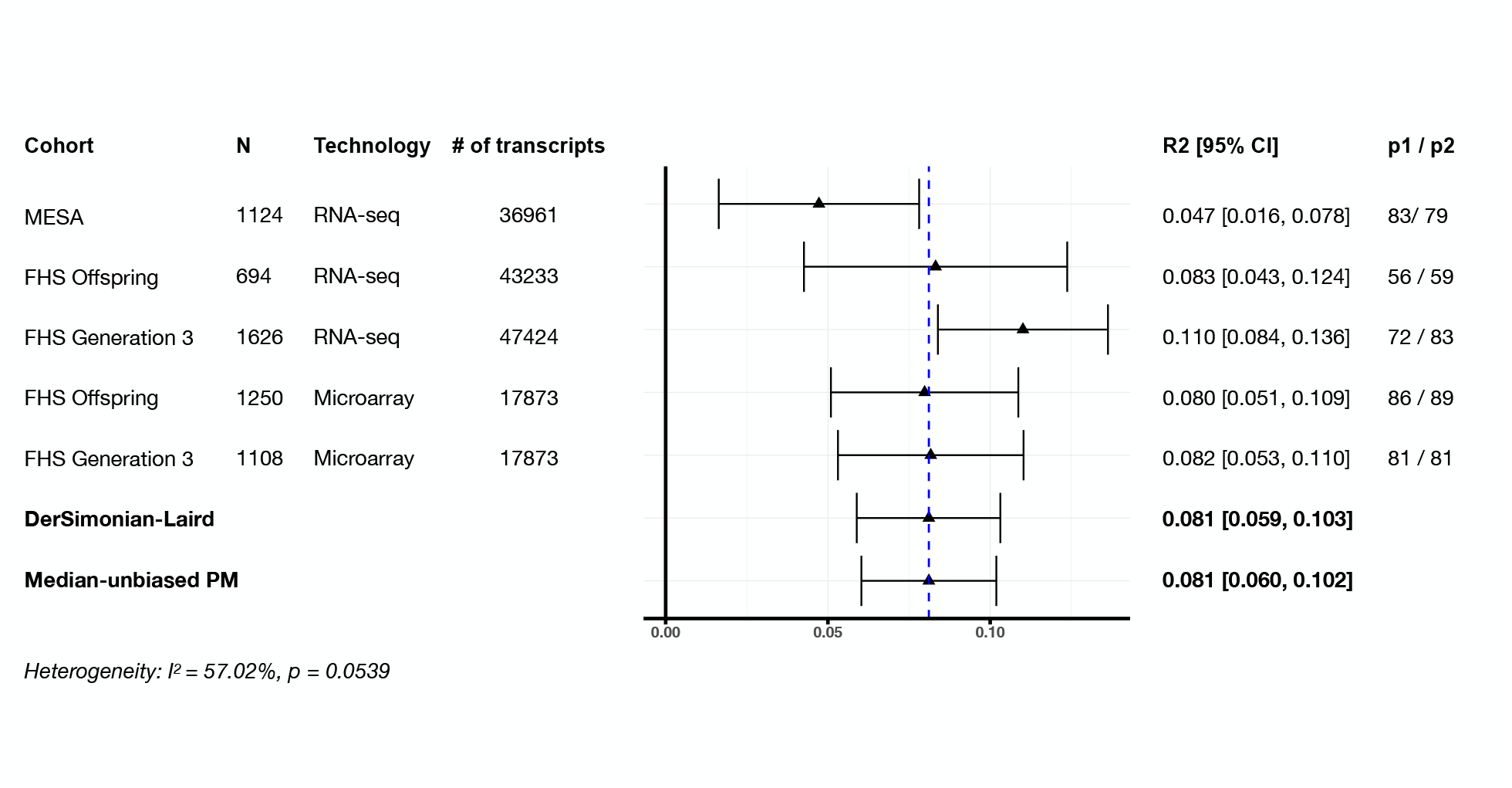
Random-effects meta-analysis results of mediation effect of gene expression between sex and HDL-C using the CF-OLS in 5 different cohorts from the NHLBI TOPMed program. *N* refers to the sample size. # of transcripts refers to the number of genes measured from the gene expression profiling. *p*1*/p*2 refers to the number of transcripts selected in the first and second subsamples, respectively. R2 refers to the total mediation effect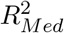. CI refers to the confidence interval.

